# Prediction of body weight from body volume of Savanna goats in Limpopo province, South Africa

**DOI:** 10.1101/2023.09.17.558151

**Authors:** Madumetja Cyril Mathapo, Thobela Louis Tyasi, Thinawanga Joseph Mugwabana

## Abstract

Savanna goats is one of the South African commercial meat breeds. This study was conducted to predict body weight from body volume (BV), heart girth (HG) and body length (BL). A total of 139 savanna goats of the age between 1 and 5 years old of different sex (male and female) were used in the study for collection of body weight, body length and heart girth. The animals were kept under semi-intensive production system, where they were supplemented in the afternoon. Body volume was derived using cylinder volume formula from body length and heart girth as the components of model. R-studio software was employed for Pearson correlation matrix to assess the association between body weight, body length, heart girth and body volume. Simple linear regression was used to establish model to predict body weight. Pearson correlation results indicated that BW had a highly statistical significant (p<0.01) correlation with HG (r = 0.90), BV (r = 0.84) and BL (r = 0.66), respectively. Regression model findings indicated that HG had highest coefficient of determination (R^2^ = 0.81) and lowest mean square error (MSE = 24.85), and BV indicated highest coefficient of determination (R^2^ = 0.76) and low mean square error of (MSE = 35.07) while BL indicated lowest coefficient determination of (R^2^ = 0.45) and high mean square error (MSE = 70.80). In conclusion, correlation result suggests that by improving HG, BV and BL will result in improving BW of Savanna goats. Simple linear regression suggest that HG and BV can be used to estimate BW of Savanna goats.

## Introduction

Savanna goats is a commercial meat breed that was developed from the indigenous goats of Southern Africa for the past decades [1]. Goats are highly fertile species that can reach maturity at an early stage with low input requirements [2]. Goat farming has been practiced in a rural level since they can survive on any available shrubs and trees in poor environment where no other crops can be found [3]. They are crucial to the livelihood of the rural farmers since they supply them with meat, milk, hides, manure and source of income [4]. Body weight is very important trait since it can direct farmers to run proper management practices in their herd and it can also be used check the development of livestock [5]. However, measuring body weight of goats is not easy at a rural level due to lack of weighing scales [6]. Then, this challenge disadvantages the farmers from running proper management of their herd and when selling since now they rely on subjective method. Estimation of body weight from morphometric traits is a cheap, easy and indirect method that can be practiced by livestock farmers at a rural level [7]. Several studies have been conducted on the estimation of body weight from various morphometric traits using different models in sheep [8], goats [9], chickens [10] and reported different traits that can be used as single predictor of body weight. However, it has been noticed that using single trait as predictor of body weight gives low precision [11]. Therefore, the use of estimating body weight from body volume in Savanna goats has not been exploited. Thus, the objective of this study is to develop a model to estimate body weight of Savanna goats from body volume (BV). Body volume is calculated from body length and heart girth of the animal. Body volume is calculated using body length and heart girth, where heart girth represents circumference of the circle cylinder shape and body length represent height cylinder of the shape [12].

## Materials and Methods Study area

The study was conducted at Game Breeder farm located in Bysteel, under Polokwane Local Municipality, South Africa. The area has an annual rainfall of 600 mm with summer temperature ranging from 16 to 28 degree Celsius and winter temperature ranging from 7 to 21 degrees Celsius [13].

### Experimental animal and management

A total of 139 goats (males and females) between the age of 1 to 5 years were used to conduct the study. The goats were kept under extensive production system, where they were allowed to go and graze and browse in the farm during the day and recall them back later into the kraal. The animals were dipped regularly since the area is known to be a heart water area. The study only focused on healthy animals, pregnant and sick animals were not included in the study.

### Data collection

Body weight was measured using a weighing scale calibrated in (kg). Body length and heart girth were measured using tailor measuring tape calibrated in (cm) following the procedure of Tesema et al. [14], where: Body length (BL) was measured as a distance between the site of pins to the tail drop while heart girth (HG) was measured as body circumference of the chest behind the forelegs. All these measurements were taken in the morning before goats were released for grazing. Body volume (BV) was calculated by cylinder volume formula calibrated in (dm^3^). Body length (BL) represented the cylinder shape and heart girth (HG) represented circumference of the circle. Therefore, the body volume (BV) in cm^3^ was calculated using the following formula:

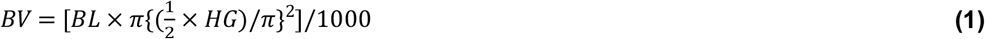

### Statistical analysis

Statistical Package for Social Sciences (IBM SPSS, 2022) version 28.0 and R-studio software was used for data analysis. Correlation matrix was used to determine the relationship between body weight, morphological traits and body volume. Simple linear regression model was used to establish a model to estimate body weight from morphological traits.

The following is a simple linear regression model that was followed:

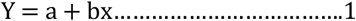

Where, Y is dependent variable (BW), a is regression intercept, b is regression coefficient and x is independent variable (BL, HG and BV).

## Results

Figure 1 shows the results of the relationship between body weight and some morphological traits. The results showed that BW had a high significant (p<0.01) correlation with HG (r = 0.90), BV (r = 0.84) and BL (r = 0.66), respectively. There was also a high significant (p< 0.01) correlation amongst the morphological traits. Where HG had high statistical significant correlation with BV (r = 0.94) and BL (r = 0.71), respectively. BV had a high significant correlation with BL (r = 0.88).

**Figure 1:**
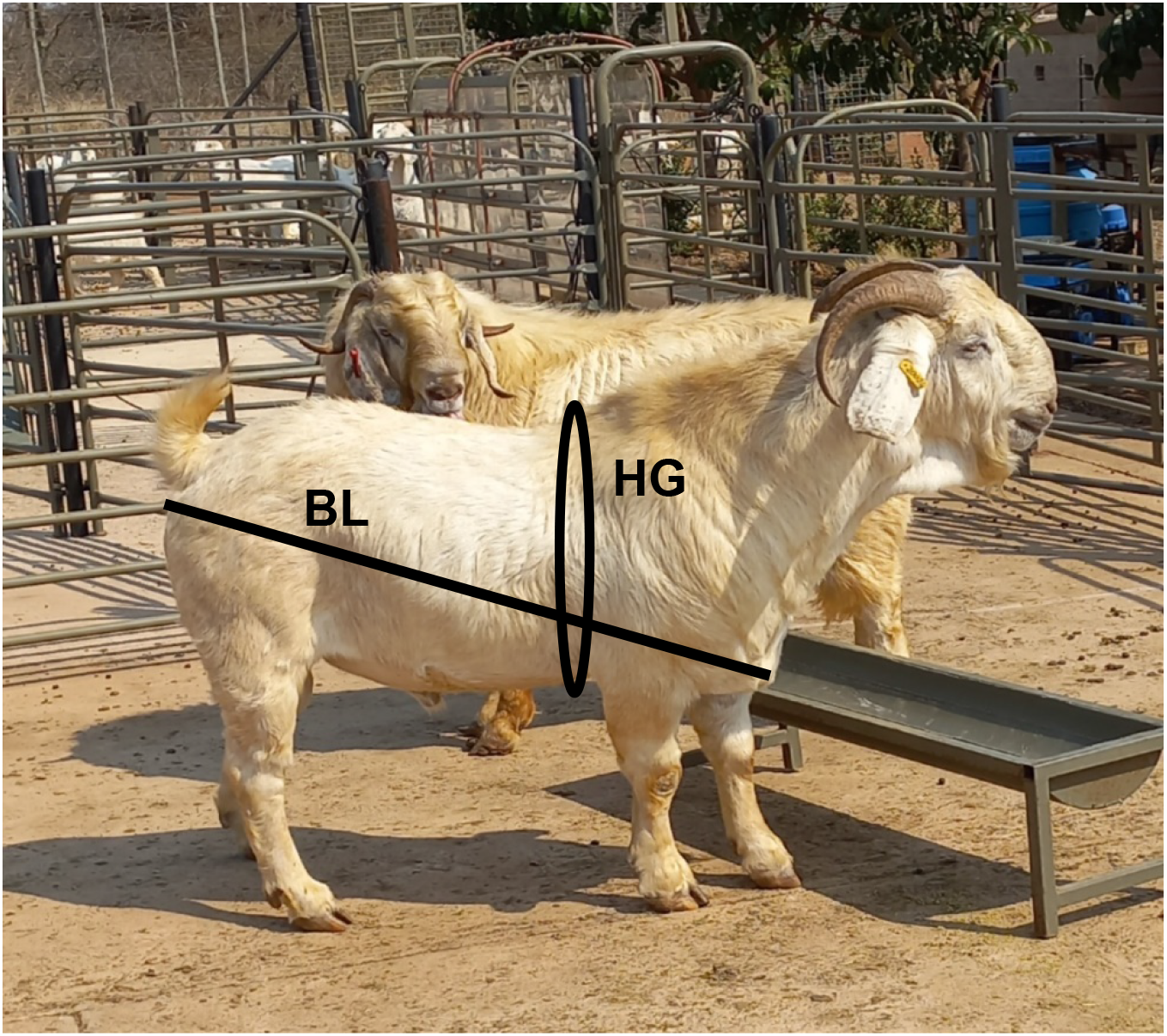
Savanna goat showing morphometric traits measured in the study.

**Figure 1:**
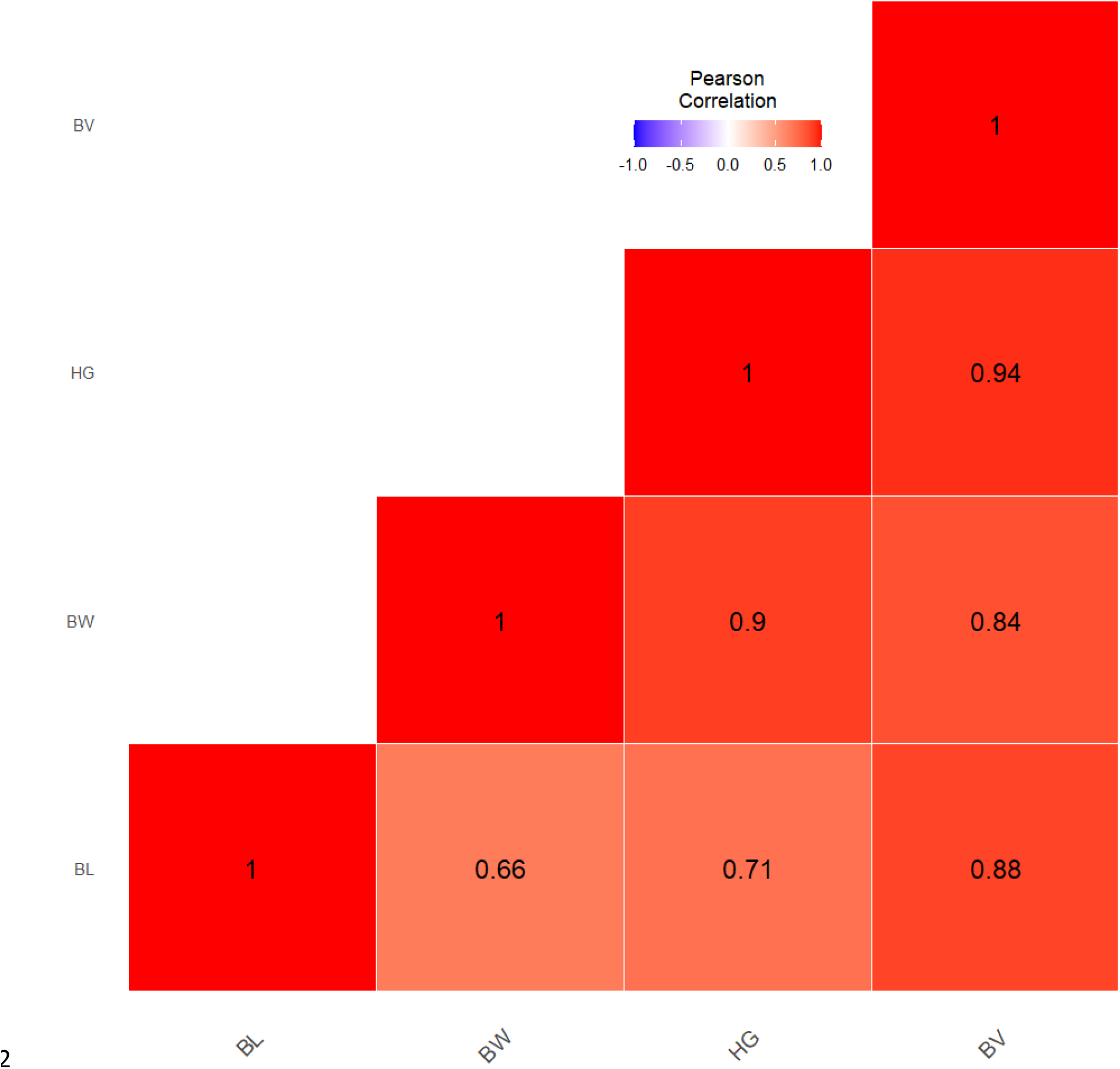
Heat map showing correlation between body weight and morphometric traits in Savanna goats. Heat map colour distribution represent the magnitude of the correlation between the variables. red colour means high correlation, blue colour low correlation and white mid correlation.

Table 1 below shows the simple linear model for prediction of body weight from morphological traits. The results indicated that HG model had the highest coefficient of determination (R^2^ = 0.90) with lowest mean square error of (MSE = 24.85), followed by BV model with coefficient of determination (R^2^ = 0.85) and mean square error of (MSE = 35.07) and BL with coefficient of (R^2^ = 0.67) and mean square error of (MSE = 70.80).

**Table 1:**
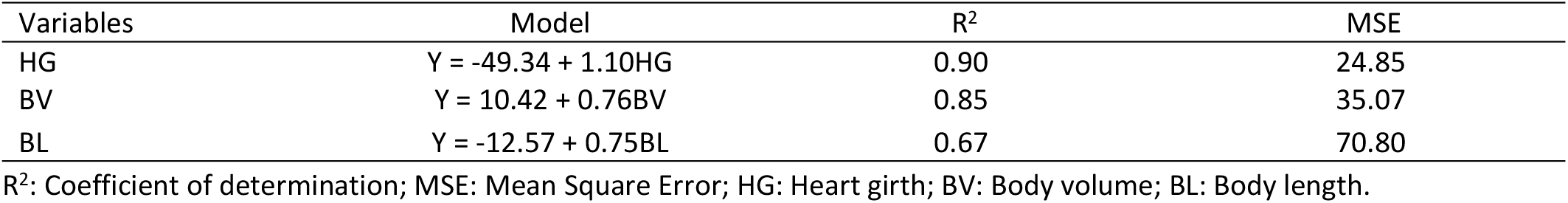
Simple linear regression model for estimation of body weight from morphological traits.

| Variables | Model | R<sup>2</sup> | MSE |
| --- | --- | --- | --- |
| HG | $Y = -49.34 + 1.10HG$ | 0.90 | 24.85 |
| BV | $Y = 10.42 + 0.76BV$ | 0.85 | 35.07 |
| BL | $Y = -12.57 + 0.75BL$ | 0.67 | 70.80 |

R<sup>2</sup>: Coefficient of determination; MSE: Mean Square Error; HG: Heart girth; BV: Body volume; BL: Body length.

## Discussion

Body volume calculated from body length and heart girth can assist in improving the precision of estimating body weight of livestock [15]. The current study firstly looked at the relationship between body weight, body volume, heart girth and body length using correlation matrix. The results of the study showed a highly significant correlation between body weight and body volume. The results of the current study are in line with the results of Gurgel et al. [16] and Salazar-Cuytun et al. [15], who showed a highly significant correlation between body weight and body volume in Santa Ines lambs and Pelibuey lambs, and sheep. The current correlation results further indicated highly statistical correlation between body weight and morphometric traits (Heart girth and body length). The results were in line with the results of other studies ([17]; [5]; [13]) who indicated a highly statistical correlation between body weight, heart girth and body length. The results means that by improving heart girth, body volume and body length will improve body weight of the Savanna goats. Thus, the study employed simple linear regression model to check the contribution of each morphometric traits and body volume to the body weight. Simple linear regression model indicated that heart girth and body volume had the highest coefficient of determination and root mean square error, respectively. The results further indicated that the body length showed low coefficient of determination and highest root mean square error. The results of the current study were in line with the results of Paputungan et al. [15] who reported highest coefficient of determination from heart girth and body volume, and low coefficient of determination of body length in Ongole crossbred cows. Takaendengan et al. [18] in horses and Salazar-Cuytun et al. [15] in hair sheep, reported results that agree with the current study results, were it was found that body volume showed highest coefficient of determination in all the models that were employed. However, low coefficient of determination and high root mean square error was observed in lactating water buffalo [19]. The result of the current study means that body volume can be used as a single predictor of body weight in the absence of weighing scale [20].

## Conclusion

This study disclosed that there is a highest association between body weight and body volume in Savanna goats. The simple linear model displayed that body volume showed a great variation on the body weight of Savanna goats. Therefore, body volume can be used to predict body weight of Savanna goats without weighing scale, due to its high accuracy. More studies need to be conducted on the prediction of body weight from body volume through different models and increase the sample size when conducting the kind of study.

## Acknowledgements

The authors are thankful to the Steven Strydom for allowing us to collect data at his farm.

